# A scalable genetic tool for the functional analysis of the signal recognition particle

**DOI:** 10.1101/2024.07.20.602779

**Authors:** Lawton F. Long, Shivani Biskunda, Ming “Peter” Yang, George C. Wu, Cassidy Simas, Steven D. Bruner, Carl A. Denard

**Affiliations:** University of Florida Department of Chemical Engineering, Gainesville, FL, 32611; University of Florida Department of Chemistry, Gainesville, FL, 32611; University of Florida Department of Biomedical Engineering, Gainesville, FL, 32611; UF Health Cancer Center, Gainesville, FL, 32611

**Keywords:** Congenital Neutropenia, Signal Recognition Particle, SRP54, *Saccharomyces cerevisiae*, Yeast disease model

## Abstract

Mutations in the *SRP54* gene are linked to the pathophysiology of severe congenital neutropenia (SCN). *SRP54* is a key protein comprising one of the six protein subunits of the signal recognition particle responsible for co-translational targeting of proteins to the ER; mutations in *SRP54* disrupt this process. Crystal structures and biochemical characterization of a few *SRP54* mutants provide insights into how *SRP54* mutations affect its function. However, to date, no scalable, flexible platform exists to study the sequence-structure-function relationships of SRP54 mutations and perform functional genomics and genome-wide association studies. In this work, we established a haploid model in *Saccharomyces cerevisiae* based on inducible gene expression that allows these relationships to be studied. We employed this model to test the function of orthologous clinical mutations to demonstrate the model’s suitability for studying SCN. Lastly, we demonstrate the suspected dominant-negative phenotypes associated with *SRP54* mutants. In doing so, we discovered for the first time that the most common yeast orthologous clinical mutation, S125del (T117del human orthologue) displayed the least severe growth defect while the less common G234E mutant (G226E human orthologue) displayed the most severe growth defect. The ability of this haploid model to recapitulate these phenotypes while remaining amenable to high-throughput screening approaches makes it a powerful tool for studying *SRP54*. Furthermore, the methodology used to create this model may also be used to study other human diseases involving essential and quasi-essential genes.

**Takeaway:**

- The constitutively expressed quasi-essential native *SRP54* gene in yeast is replaced with an aldosterone-inducible version, making a yeast haploid strain, LFL001, that requires aldosterone for growth.
- Strains developed in this work corroborated that yeast *SRP54* mutations exhibit dominant-negative phenotypes.
- The LFL000 and LFL001 yeast strains developed here are uniquely positioned to interrogate sequence-structure-function relationships of *SRP54*.

## Introduction

Congenital neutropenias (CNs), which include severe congenital neutropenia (SCN) and Schwachman-Diamond-like syndrome (SDS), are a class of rare, uncurable bone marrow failure diseases characterized by low neutrophil levels (Thompson et al., 2022). Children with CN show high susceptibility to infections, various organ dysfunctions, and present a high risk for leukemic transformation (Sabulski et al., 2022; Touw & Beekman, 2013). While over 20 genes associated with CN have been discovered, heterozygous mutations in the signal recognition particle 54 gene (*SRP54*) have recently been identified as a cause of SCN and SDS (Bellanné-Chantelot et al., 2018; Kellogg et al., 2022), suggesting that protein co-translational targeting plays a major role in CN. The signal recognition particle (SRP) promotes co-translational targeting of a nascent protein bearing an N-terminal signal peptide to the endoplasmic reticulum (ER) (Arnold et al., 1998; Hann & Walter, n.d.) (Brown et al., 1994). The eukaryotic SRP is composed of 6 protein subunits (*SRP9*, *SRP14*, *SRP19*, *SRP54*, *SRP68*, and *SRP72*) and a 7SL RNA core (Juaire et al., 2021; Walter & Blobel, 1980). The *SRP54* subunit is responsible for binding the nascent signal peptide emerging from the ribosome while its GTP-driven binding with the *SR*/11 (an SRP receptor protein) facilitates successful protein docking with *SR*β (an SRP docking protein), resulting in protein translocation into the ER.

In recent years, several studies have connected mutations within *SRP54* to SCN and SDS (Bellanné-Chantelot et al., 2018; Sabulski et al., 2022). *SRP54* comprises two domains: the NG domain responsible for GTP binding and a methionine-rich (M-domain) responsible for signal peptide recognition. CN stemming from *SRP54* mutations occurs via mutations in the NG domain (T115A, G226E, and T117del). These mutations impact GTP binding resulting a loss of SRP function (Juaire et al., 2021). Studies conducted in zebrafish have also demonstrated that mutations within *SRP54* may result in dominant-negative phenotypes (Schürch et al., 2021). While these findings suggest that *SRP54* mutations involve complex molecular interactions, there is a lack of existing platforms for further research into these molecular mechanisms (Schürch et al., 2021). SCN is estimated to have an occurrence rate between 3 and 8.5 cases per million people globally (Skokowa et al., 2017). While mutations in the *ELANE* and *HAX1* genes have been known, only recently have SRP54 germline mutations been identified as notable contributors to SCN. Although based on few reports, *SRP54* mutations are the second most common cause of CN with maturation arrest and comprises 6.9% in the French CN registry (Bellanné-Chantelot et al., 2018). Therefore, efforts to invigorate research into *SRP54* and its relationship to SCN would benefit from a scalable genetic tool that facilitates functional studies.

The yeast *Saccharomyces cerevisiae* is a stalwart chassis to study human diseases that involve protein secretion, proteostasis, and cell division (Bolotin-Fukuhara et al., 2010; Kachroo et al., 2022). Since the SRP complex is highly conserved across all domains of life (Brown et al., 1994), *S. cerevisiae* can model SRP function and its associated molecular processes (Keenan et al., 2001; Sheara & Jeffrey, n.d.). In this work, we developed a scalable and genetically tractable haploid *S. cerevisiae* platform to study the impact *SRP54* mutations have on overall cellular function and to gain insight into the underlying mechanisms of SCN. In this model, the expression of *SRP54* can be toggled “on” or “off” and titrated to induce disease phenotypes without impacting cell viability during non-experimental steps. Importantly, our genetic tool does not rely on isolating haploids from engineered diploids. Our SRP54 yeast models will allow us to directly study sequence-structure-function relationships (Fowler & Fields, 2014; Ravikumar et al., 2018), conduct cost-effective functional genomics studies and high-throughput drug screening.

## Results

*SRP54* is a quasi-essential gene whose deletion in *S. cerevisiae* causes a severe growth defect (Brown et al., 1994; Juaire et al., 2021). Conventional approaches to manipulate essential genes include temperature-sensitive haploids and the use of diploids (Giaever & Nislow, 2014; Kofoed et al., 2015). However, temperature-sensitive mutants are experimentally restrictive and cannot support genetic circuits with fast response times. Diploids, including a recent hormone-inducible collection (Arita et al., 2021), retain a chromosomal copy of the WT gene, making them unsuitable for uncovering the direct impact of mutations on function. Moreover, tetrad dissection (Ravikumar et al., 2018; Wellner et al., 2021) is not conducive to high throughput methodologies (Morin et al., 2009). Thus, we envisioned a haploid yeast strain where *SRP54* expression is driven by an inducible, titratable promoter. Titratable “turn on” transcription of *SRP54* supports normal cell growth in preparation for large-scale functional studies. Contrariwise, functional genomics experiments, including cellular rescue and phenotypic and deep mutational scanning assays, would occur without an inducer of *SRP54* expression. Similar approaches that replace genes with galactose-inducible counterparts, which include cases for *SRP54*, impose a major metabolic shift within the organism (Mutka & Walter, 2001; Sanford et al., 2022), resulting in slower induction kinetics and inability to titrate gene expression.

We aimed to build a scalable, controllable genetic tool that would enable sequence-structure-function studies of *SRP54*. Our model strain contains a deletion of the native *SRP54* deletion but harbors a surrogate copy of the gene under an aldosterone-inducible promoter. In this approach, cell growth and viability could be decoupled from functional studies (Figure 1). In addition, this approach leverages tunable gene expression capabilities afforded by hormone-inducible systems while maintaining the engineering flexibility offered by a haploid strain. In our scheme, addition of aldosterone to the cell culture media toggles *SRP54* gene expression “on” resulting restored cell growth (LFL001). The gene expression can be toggled “off” by growing cells in media lacking aldosterone resulting in a growth defect.

**Figure 1:**
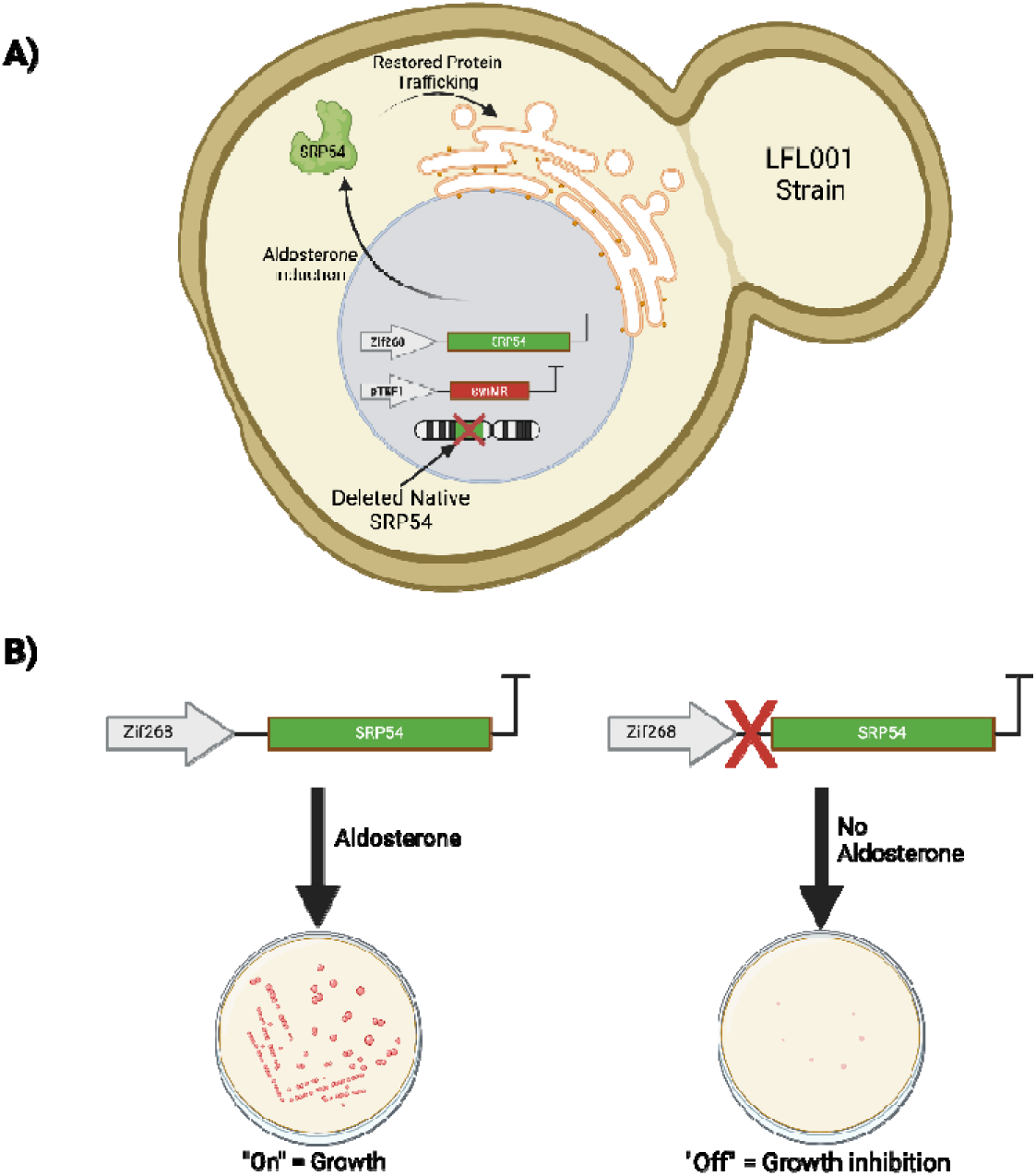
Features of yeast SCN model. **A)** Depiction of LFL001 strain when induced with aldosterone. Addition of aldosterone restores ER-dependent protein trafficking allowing for cell growth in *SRP54* deletion strain. Removal of aldosterone results in reversion to growth-inhibited phenotype in *SRP54* deletion strain. **B)** Phenotype exhibited in absence of (Left) and in the presence of (Right) aldosterone.

### Generating the LFL001 strain

To generate LFL001, we first built two separate integration vectors. One harbors a synthetic transcription factor, *synMR*, under the control of the strong constitutive promoter *pTEF1*; *SynMR* is required to drive gene expression with aldosterone addition (Sanford et al., 2022). The second vector contains an *SRP54* gene driven by an aldosterone-inducible promoter, *pMR*. The *pMR* promoter has a high activation threshold compared to other synthetic promoters and can be titrated to reach a level of maximal gene expression equivalent to the galactose-inducible promoter (Sanford et al., 2022). This high activation threshold makes the promoter less prone to leaky expression. To minimize the impact of codon usage on protein expression level, we kept the DNA sequence for the aldosterone driven *SRP54* (called *SRP54*\* from here on) the same as the native yeast *SRP54* with only minor differences. Specifically, we made 9 silent mutations in the M domain of the aldosterone driven *SRP54*\*. Anticipating that CRISPR-Cas would be used to delete the host cell’s *SRP54* gene, we thus ensure that guide RNAs that target the host *SRP54* would not impact *SRP54*\* (Supplemental Figure 1) (Laughery & Wyrick, 2019).

With the transcription factor and aldosterone-inducible *SRP54*\* cassettes integrated into the yeast chromosome (we call this first strain LFL000), we deleted the native *SRP54* to create the final strain (LFL001) (Supplemental Figure 2) using a markerless deletion approach. To do this, we designed a guide RNA (gRNA) that targets the M domain of the native *SRP54* (Laughery & Wyrick, 2019). This gRNA is thus unique to the native *SRP54* gene thanks to the silent mutations introduced in the *SRP54\** M-domain. Additionally, we designed a 200-base pair markerless repair template with homology arms flanking the *SRP54* gene, leading to the deletion of the entire *SRP54* coding sequence. The native *SRP54* is driven by a bidirectional promoter that also transcribes a gene of unknown function; therefore, the repair template ensured that the *SRP54* promoter (and terminator) remained unchanged (Supplemental Figure 3). To delete the native *SRP54* gene, cells were transformed with the Cas9 + gRNA vector and plated on SC-Leu + 10 µM aldosterone to select for the Cas9 plasmid and deletion while turning on *SRP54*\* gene expression to maintain cell growth during this stage. In a subsequent step, colonies were screened for the expected growth defect by streaking each colony on a YPD plate with and without 10 µM aldosterone. A colony with the expected growth defect was isolated and genotyped to confirm the presence of the 200-base pair repair template (Figure 2).

**Figure 2:**
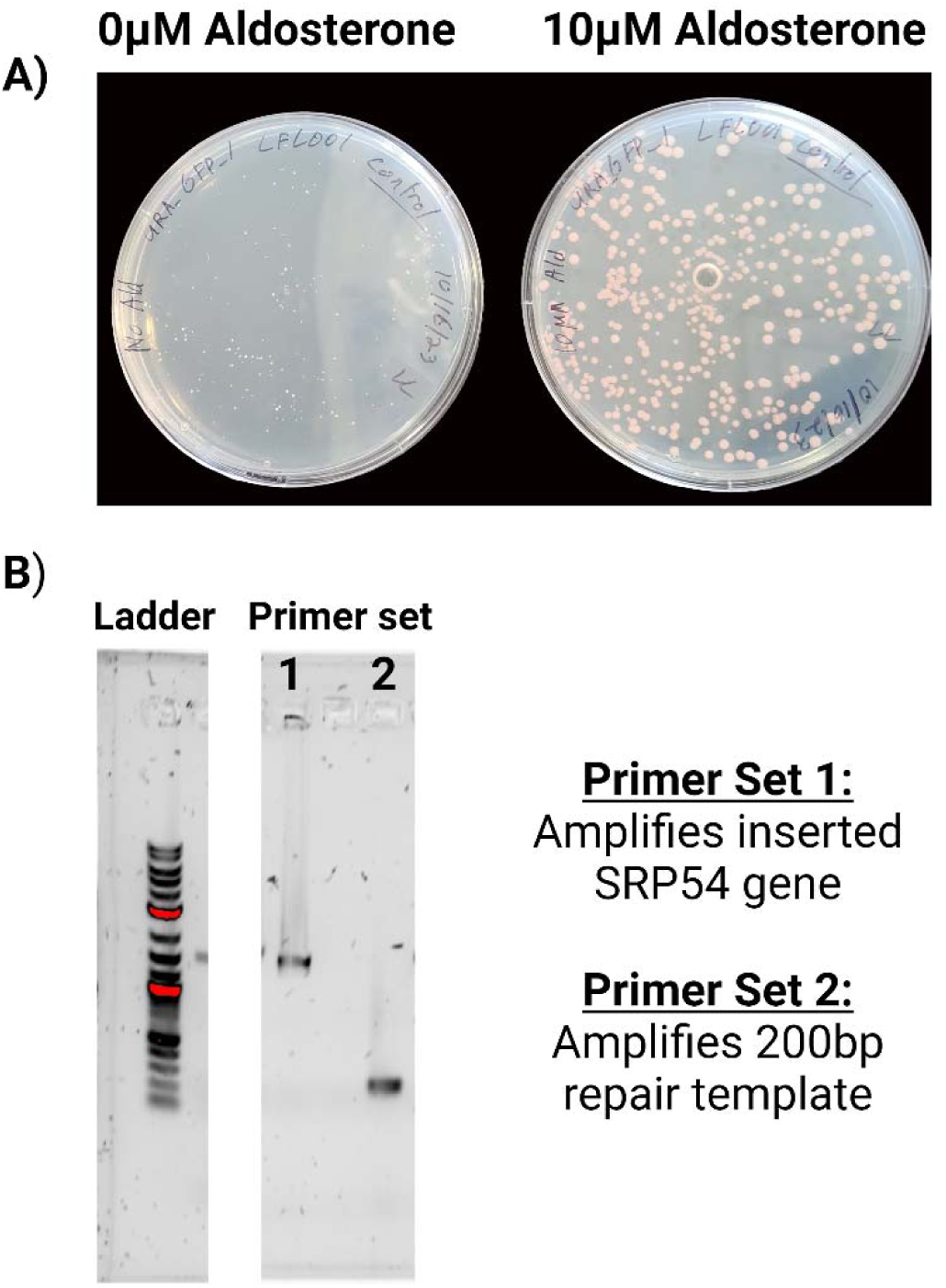
Genotyping of LFL001 strain. **A)** LFL001 growth in the presence of 10 µM aldosterone (right) and in the absence of aldosterone (left) after 3 days at 30C. Samples treated with aldosterone exhibit significantly faster growth than samples without aldosterone. **B)** Primer set 1 indicates PCR amplification of *pMR*-*SRP54*\* cassette from genomic DNA isolated after knockout of the native *SRP54*. Primer set 2 verifies deletion of the native *SRP54*.

### LFL001 growth is dependent upon aldosterone induction

The growth of LFL001 in liquid media was compared to that of the starting strain, W303-1a, with the *SRP54*\* turned “on” and “off” (Figure 3). At an aldosterone concentration of 0.625 µM, *SRP54*\* expression was high enough to restore growth significantly and LFL001 reached a maximum OD_600_ of 15. In contrast, only slight growth was observed in the absence of aldosterone in the culture media at early time points < 24 hours, with the LFL001 strain reaching an OD_600_ of approximately 5 at ∼40 hours before growth stagnation (Figure 3). These findings are consistent with prior research showing that *SRP54* is quasi-essential for cell growth (Brown et al., 1994). Higher aldosterone concentrations (2.5µM, 5µM, and 10µM) did not further increase cell growth rates. Despite a successful growth recovery, the W303-1a strain still grew at a faster rate than the LFL001 strain and reached the expected maximal OD_600_ of 20. This difference in growth rate and final density suggests that epigenetic and DNA context-dependent mechanisms may regulate the SRP pathway within *SRP54* (Mayr, 2019). The ability to tune *SRP54*\* expression levels to match WT *SRP54* gene expression is a key advantage of LFL001. RT-qPCR reveals that *SRP54* expression levels in LFL001 closely match native *SRP54* gene expression levels at 0.625µM aldosterone but becomes overexpressed at concentrations greater than 2.5µM (Supplementary Figure 4). Taken together, these experiments show that the design of LFL001 strain was successful and that LFL001 could be used as a model for *SRP54* functional studies.

**Figure 3:**
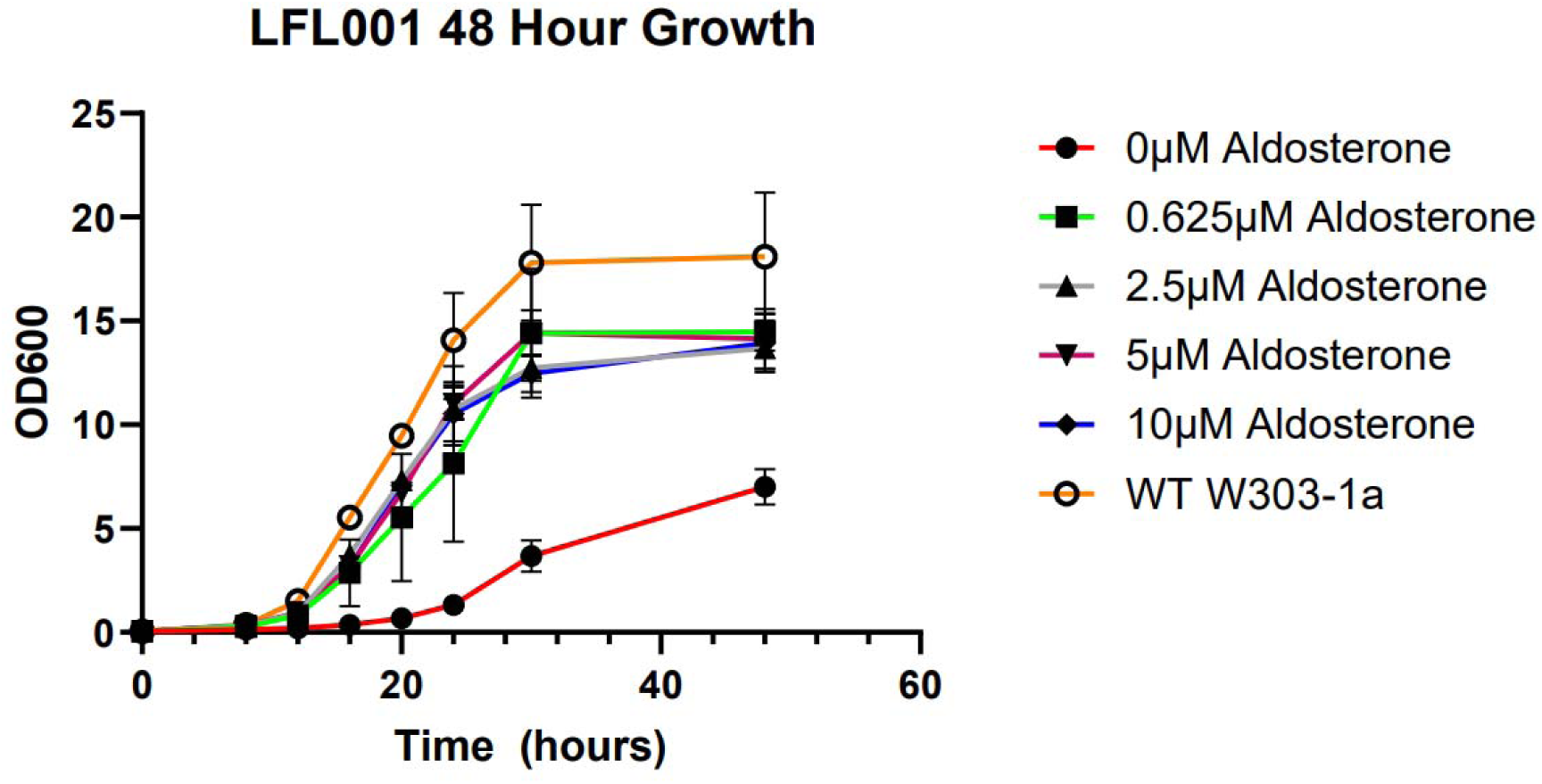
LFL001 is dependent on aldosterone for growth. Growth characteristics of LFL001 strain compared to the base W303-1a strain. LFL001 displays slightly slower growth compared to WT W303-1a but is significantly faster than the 0 µM control. Aldosterone concentration does not seem to significantly affect the growth rate of the LFL001 strain at concentrations > 0.625 µM. LFL001 reaches a lower final OD_600_ than the base W303-1a strain used to create the LFL001 strain.

### LFL001 can replicate loss of function phenotypes resulting from *SRP54* mutations

To characterize loss of function *SRP54* mutations, 3 mutations orthologous to common human clinical mutations were tested: S125del, T123A, and G234E (orthologous mutations in human SRP54 are T117del, T115A, and G226E, respectively) (Juaire et al., 2021). These mutants are unable to complement growth in a previously built *SRP54* deletion haploid strain (Juaire et al., 2021). To verify these growth defective phenotypes, we placed the mutants on a plasmid under the control of the *pADH1* promoter and transformed them into the LFL001 strain (Figure 4). For rescue experiments, transformed LFL001 cells were plated on media without aldosterone. Expectedly, none of the 3 mutants could complement cell growth without aldosterone addition (Figure 4).

**Figure 4:**
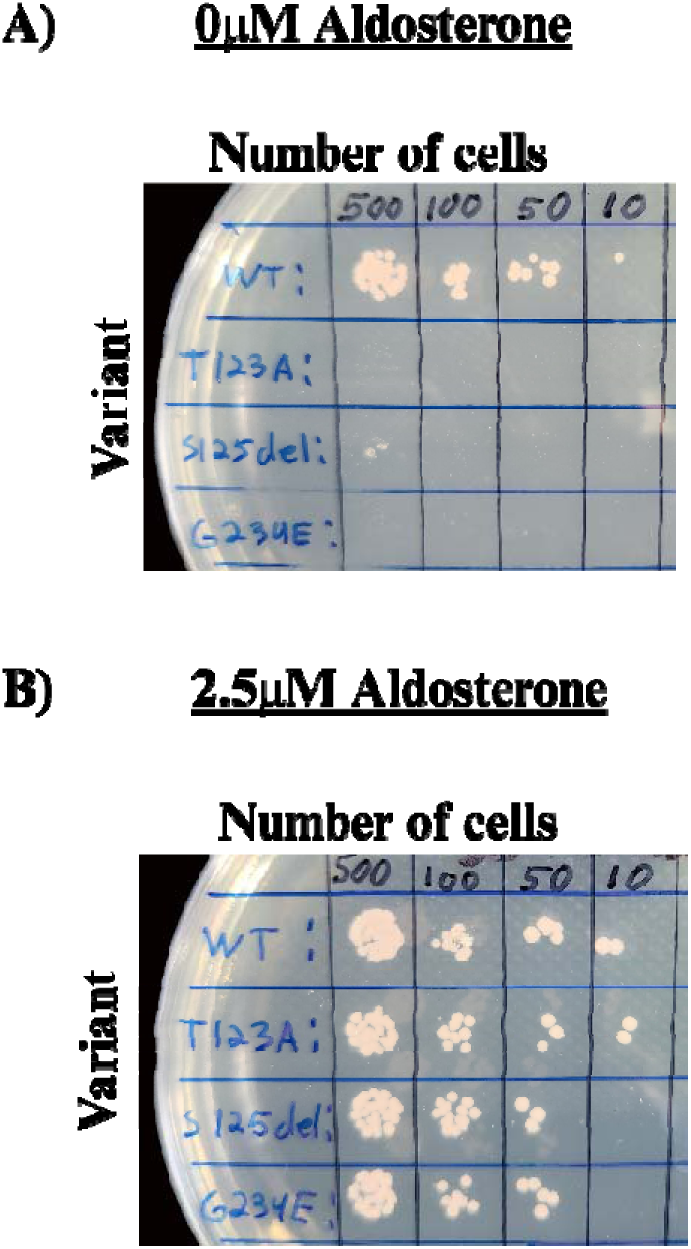
Yeast mutant rescue assay in LFL001. *pADH1*-driven *SRP54* mutants cannot complement cell growth. **A)** WT, T123A, S125del, and G234E variants were transformed into LFL001 and spotted on an SC-Ura plate with no aldosterone. All 3 mutants exhibited severe growth defects after 3 days. **B)** The variants were then spotted on an SC-Ura plate containing 2.5µM aldosterone as a positive control. No significant difference in colony size was observed indicating that dominant-negative phenotypes were unable to be observed in this configuration. It is believed that this lack of observed difference between the samples was due to WT expression levels being significantly higher than that of the expressed mutants.

### Yeast *SRP54* mutations exhibit dominant-negative (DN) phenotypes

Clinical mutations in human *SRP54* exhibit dominant-negative phenotypes in Zebrafish; however, whether and to what extent these phenotypes exist in the yeast orthologue has not been systematically examined (Juaire et al., 2021; Schürch et al., 2021). To determine whether these dominant-negative phenotypes occur in yeast, we inserted the 3 mutants into a LFL000. LFL000 is a W303-1a strain that harbors the *synMR* transcription factor required to drive the *pMR* promoter but retains a WT copy of *SRP54* under its native promoter. The mutants were placed on a plasmid under control of the *pMR* promoter allowing their expression to be titrated with varying aldosterone concentrations. This configuration enables us to vary the transcriptional stoichiometric ratio of the mutant *SRP54*. Importantly, matching the expression levels of the mutant to that of the WT provides a more accurate representation of dominant-negative phenotypes in human cells (Birchler & Veitia, 2010). Indeed, Figure 5 shows that *SRP54* mutant toxicities emerge at 1.25 µM aldosterone and are exacerbated by increased aldosterone concentrations. From these results, one can summarize that the T123A mutant demonstrated a moderate dominant-negative phenotype. In contrast, the S125del mutant displayed the least severe dominant-negative phenotype. Interestingly, G234E mutant exhibited the most severe dominant-negative phenotype. GTP binding assays in the human orthologue of G234E, G226E, show a two-fold increase in GTP binding affinity (Juaire et al., 2021). We hypothesize that the yeast G234E mutant also has increased GTP binding affinity compared to WT *SRP54* (Juaire et al., 2021). This retained GTP binding ability would allow the G234E mutant to bind to other subunits of the SRP complex, such as *SR*/11, interfering with WT function. These dominant-negative phenotypes arise from the ability of the respective mutants to effectively poison the SRP complex (Herskowitz, 1987; Veitia, 2008).

**Figure 5:**
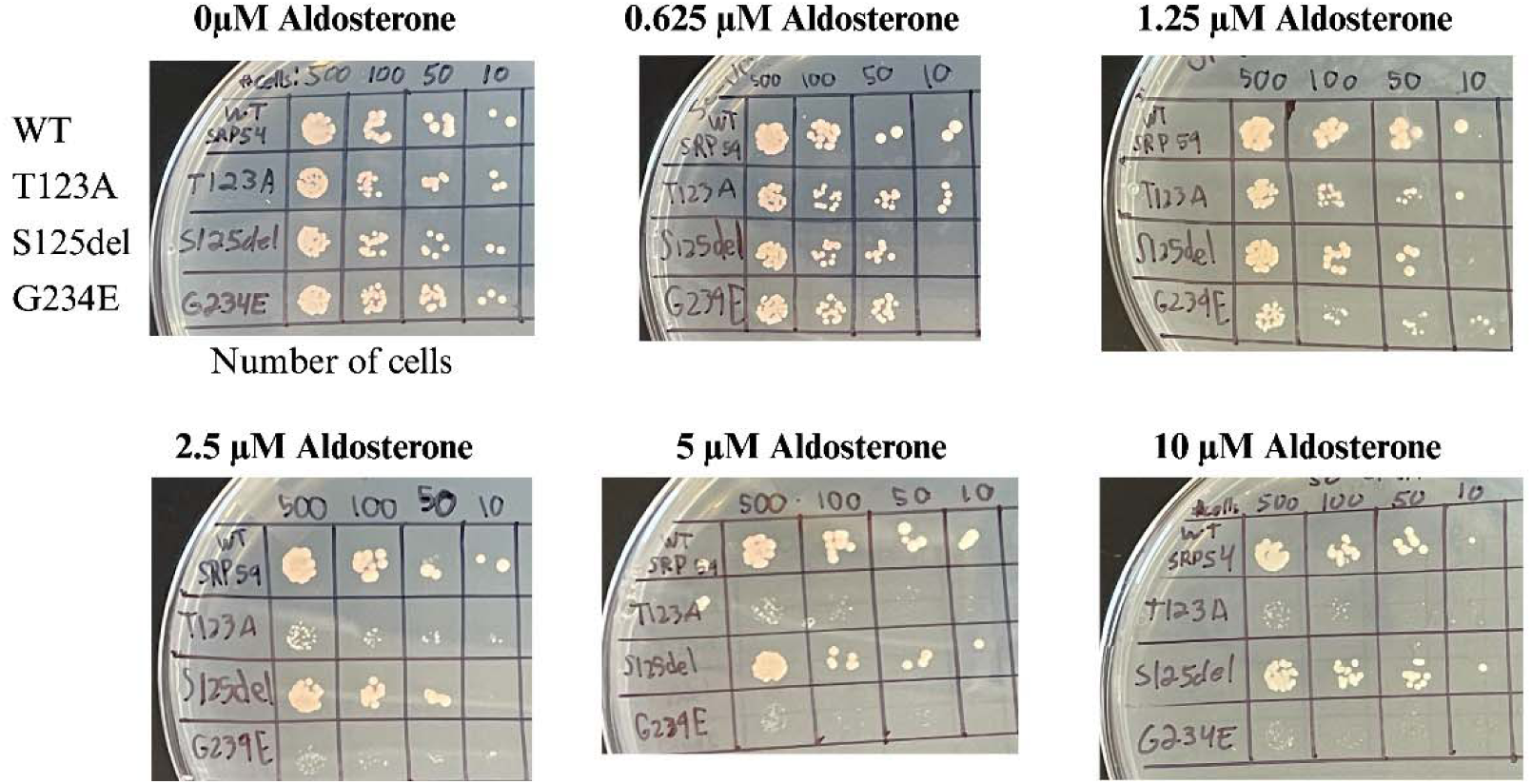
Demonstration of dominant-negative mutant phenotypes in LFL000. Spotting assay in the LFL000 strain where chromosomal WT *SRP54* is retained, and a *pMR*-driven mutant is inserted into a plasmid. The aldosterone concentration was increased from 0.625 µM to 10µM to titrate the dominant-negative phenotype for each mutant. The dominant-negative phenotype for the G234E and T123A mutants became observable at 1.25 µM with the G234E displaying the most severe growth defect. The S125del mutant was the least severe and did not exhibit any discernible dominant-negative phenotype.

## Discussion

In this work, we demonstrated the utility of a haploid model of *SRP54* (LFL001) that leverages a hormone inducible and titratable approach to control gene expression. We established the reliability of this strain by characterizing three yeast *SRP54* mutants (T123A, S125del, and G234E) which are orthologous to clinical mutations in human *SRP54* associated with SCN. As expected, none of the 3 mutants were able to complement cell growth. Additionally, our LFL000 strain could accurately recapitulate dominant-negative phenotypes bought upon by *SRP54* mutations in a diploid-like setup. We find that the SRP54 mutations display varying levels of dominant-negative toxicities. The S125del mutant (orthologous to human T117del) displays little toxicity in this model, while the T123A mutant (orthologous to human T115A) displays moderate toxicity. Notably, we demonstrate that the G234E (orthologous to human G226E) mutant expression results in a severe dominant-negative phenotype. These findings are in line with experiments in zebrafish where mRNA injection of the T115A human *SRP54* variants exhibit more severe phenotypes than the T117del variant (Carapito et al., 2017; Schürch et al., 2021). Importantly, the G226E mutation found in patients with SCN has been demonstrated to have two-fold higher GTP binding affinity than WT which may prevent turnover of GTP after hydrolysis. Increased GTP binding affinity may also be responsible for the significant dominant-negative phenotype observed with the G234E yeast orthologue. By comparison, the T123A yeast mutant displays a moderate dominant-negative phenotype. Its human orthologue, T115A, retains GTP binding affinity, albeit lower than WT. Lastly, the T117del human SRP54 mutant (ortholog in yeast is S125del) loses its ability to bind GTP, which in turn, leads to the least severe form of dominant-negative toxicity. Taken together, these observations suggest that variants with retained GTP binding ability are more disruptive than those without (Juaire et al., 2021).

The methodology used to create the LFL001 strain can be applied to other quasi-essential and essential genes. Because models created in this manner always retain a functional copy of the gene, deletion of the native gene should not result in lethality. Creation of essential gene models in this manner provides an advantage over traditional methods used to create haploids from diploids that rely on sporulation and tetrad dissection which are not conducive to high throughput genetic studies. By directly engineering a haploid parent, our strain is primed for large scale functional genomics studies afforded by the wide array of yeast synthetic biology tools. Therefore, haploid models like LFL001 provide unique utility for studying sequence-structure-function relationships for essential and quasi-essential genes (Arita et al., 2021; Giaever & Nislow, 2014).

## Materials and methods

### Cloning and plasmid assembly

In our design scheme, *SRP54*\* isintegrated in the yeast chromosome prior to deleting the native *SRP54* (Lee et al., 2015). The *pTEF1*-driven *SynMR* TF and the *pMR* driven *SRP54*\* were sequentially integrated in the chromosome of S. cerevisiae W303-1, with the *SynMR* at the *HO* locus under nourseothricin selection (NatR), and *SRP54*\* integrated at the *Met17* locus under hygromycin B selection. The *pMR* promoter and *SRP54*\* were assembled into an integration vector with a hygromycin B resistance marker (HygR) using the MoClo-YTK for integration at the *Met17* locus (Sanford et al., 2022). With the 2 integration vectors created, a series of integrations were done to integrate both cassettes at their respective loci in the yeast genome. These integrations were done by linearizing the respective integration vectors with the NotI restriction endonuclease to expose the chromosomal homology regions required for successful integration (Lee et al., 2015).

All plasmids used for this study are listed in Supplementary Table 1. Unless otherwise mentioned all plasmids were assembled using the MoClo-YTK toolkit using Golden Gate cloning techniques (Lee et al., 2015). This kit was a gift from John Dueber (Addgene kit #1000000061). Both the *pMR* promoter used to drive *SRP54*\* gene expression as well as the *synMR* transcription factors were gifts from Ahmad Khalil (Addgene #194491 and #194481 respectively). Site-directed mutagenesis was used to create all mutants used in this work via *HiFi* assembly using complementary primers and NEBuilder *HiFi* DNA assembly master mix (NEB Cat # E2621S) (Olszakier & Berlin, 2022). The yeast *SRP54* used to create all DNA constructs for this work was purchased from Twist Bioscience on a chloramphenicol-resistant *E. coli* vector. The gene was flanked on both ends by external BsaI restriction enzyme sites such that the resulting overhangs were compatible with MoClo-YTK promoters and terminators (Lee et al., 2015; Sanford et al., 2022). All oligonucleotides were purchased from Sigma-Aldrich. Transformations into *E. coli* were performed using NEB Turbo (Cat # C2984H) and NEB 5-alpha (Cat # C2987H) strains. All enzymes used were purchased from NEB.

### Yeast strains and media

Yeast strains used for this work are listed in Supplementary Table 2. Aldosterone used was purchased from Cayman Chemical (Cat # 15273) and was dissolved in DMSO to a concentration of 10mM. Yeast dropout media supplements were purchased from Sunrise Science. Most yeast culturing was done using synthetic culturing media (1.92g/L dropout media supplement, 6.7 g/L yeast nitrogen base w/o amino acids (Becton Dickinson), 20g/L alpha-D-glucose (Sigma Aldrich). YPD media was used when selection was not required (20g/L bacto peptone, 10g/L yeast extract, 20g/L alpha-D-glucose). When solid growth media was required, 20g/L of bacto agar (BD Difco) was added in addition to the media components.

### Yeast transformations and integrations

Competent cells used for all plasmid-based yeast transformations were prepared using the Zymo Research Frozen EZ-Yeast Transformation II kit (Cat # T2001). Unless otherwise stated all transformations were done using 200ng of plasmid and 50uL of cells and the transformation protocol outlined in the Zymo Research Frozen EZ-Yeast transformation II kit. Note that to prepare competent cells for DNA transformation, LFL001 cells were grown in the presence of aldosterone. Yeast integrations were performed according to a protocol adapted from *Geitz and Woods* (Daniel Gietz & Woods, 2002). In this method, competent cells were prepared the day of integration starting at an OD of 0.2. The cultures were grown at 30 °C in an incubating shaker at 250rpm to a final OD_600_ between 0.8 and 1.4. The cells were then transferred to a 15mL conical tube and centrifuged at 3000g for 5 minutes at room temp. The supernatant was removed, and the cells were washed in 5mL sterile water and centrifuged again at 3000g for 5 minutes. The supernatant was removed, and the cells were resuspended in 1mL sterile deionized water and the cells were centrifuged at 3000g for 5 minutes. Cells were aliquoted in 50uL aliquots and were spun at 13,000 rpm for 30 seconds in a tabletop centrifuge and the supernatant was removed. Cells were combined with the following reagents: 260µL PEG 3350 (50% W/V), 36µL LiOAc (1.0M), 10µL Salmon Sperm DNA (10µg/µL), 54µL DNA (5µg) + H2O. The cells were then incubated at 42 °C for 40 minutes. For auxotrophic markers, the 200uL of cells were plated on prewarmed plates containing the appropriate selection media and incubated at 30 °C for 3 to 5 days. For antibiotic selection, cells were rescued for 2 hours in 1mL YPD media at 30°C in a culture tube spun down at 3000g and the pellet resuspended in 200µL sterile water before plating.

### Yeast spotting assays

Yeast spotting assays were conducted using SC-Ura dropout media. 2mL liquid cultures were inoculated using a colony isolated from a re-streak of the respective strain 24 hours prior to each assay in SC-Ura media. Cultures containing LFL001-based strains were grown in 1uM aldosterone and strains in the LFL000 strain were grown without aldosterone. Prior to preparing the dilutions used in the spotting assay 1mL of each culture was first washed in 50mL TE buffer (pH 8.0) prior to resuspending in 1mL sterile deionized water. Dilutions were prepared such that each 5µL spot was estimated to contain 500, 100, 50, and 10 cells respectively. These calculations were made using the assumption that each culture contained approximately 1.5 x 10^7^ cells per OD_600_ unit per 1mL of culture. Dilutions were prepared in a 96 well plate by adding the required culture volume to achieve the respective dilutions to 150µL of sterile deionized water. Prior to incubation at 30 □ each plate was left at room temperature with the agar side up to allow each 5µL spot time to absorb into the agar plate. This was done to mitigate the risk of disturbing the spots prior to transportation to the incubator where they were incubated upside down at 30 □ for 3 days.

### Quantification of gene expression by qRT-PCR

To quantify gene expression LFL001 was grown in YPD at various aldosterone concentrations: 0µM, 0.625µM, 2.5µM, 5µM, and 10µM. In parallel, WT W303-1a was grown in YPD. RNA was harvested from the samples after 13 hours of growth at 30L. Quantification of gene expression was conducted using qPCR and analysis was performed using the ΔΔCt. RNA was harvested from all yeast cultures using a Zymo Research YeaStar RNA kit (Cat #R1002). All qPCR was conducted using a Bio-Rad iTaq Universal SYBR Green One-Step kit (Cat# 1725150). The machine used for qPCR was a Bio-Rad CFX96 Touch Real-Time PCR Detection System.

## Acknowledgements

We thank other members of the Denard Lab Group for their thoughtful discussion and input on this work. We also thank the Bruner Lab members for their suggestions regarding experimental design and direction.

