## Supplemental File for "A scalable genetic tool for the functional analysis of the signal recognition particle"

**Supplemental Table 1:** List of relevant DNA constructs used for this study

| **Plasmid Name** | **Description** |
| --- | --- |
| Met17_Zif268_SRP54 | Integration vector used to integrate *pMR*-*SRP54* at the *Met17* Locus. |
| HO_NatR_synMR | Integration vector used to integrate the s*ynMR* transcription factors at the *HO* locus. |
| URA3_SRP54WT | p*ADH1*-driven WT *SRP54* used for rescue assay. |
| URA3_SRP54_T123A | p*ADH1*-driven T123A *SRP54* variant used for rescue assay. |
| URA3_SRP54_S125del | p*ADH1*-driven S125del *SRP54* variant used for rescue assay. |
| URA3_SRP54_G234E | p*ADH1*-driven G234E *SRP54* variant used for rescue assay. |
| Zif268_URA3_SRP54WT | *pMR*-driven *SRP54* used for rescue assay. |
| Zif268_URA3_T123A_SRP54 | *pMR*-driven T123A *SRP54* variant used to test for dominant-negative phenotype. |
| Zif268_URA3_S125del_SRP54 | *pMR*-driven S125del *SRP54* variant used to test for dominant-negative phenotype. |
| Zif268_URA3_G234E_SRP54 | *pMR*-driven G234E *SRP54* variant used to test for dominant-negative phenotype. |
| Twist_ySRP54_mutCutsite | Twist *SRP54* (YTK part 3) plasmid compatible with MoClo-YTK assembly via BsaI golden gate assembly. |
| PML107_SRP54_gRNA_2 | Cas9/sgRNA expression vector used to delete the native *SRP54* gene. |
| SRP54 HDR template | Repair template used to delete the native *SRP54* gene via Cas9-mediated homology directed repair. |

Benchling link: <https://benchling.com/lawtonfl3/f_/80BFqhY9-srp54-paper-plasmids/>

**Supplemental Table 2:** List of yeast strains and genotypes

| **Strain name** | **Description** | **Genotype** |
| --- | --- | --- |
| W303-1a | Base strain used to create LFL001 and LFL000. | MATa ade2-1 ura3-1 his3-11 trp1-1 leu2-3 leu2-112 can1-100 |
| LFL000 | Strain retaining a native SRP54 copy and the required *synMR* transcription factors to drive the *pMR* promoter. | MATa HO::pTEF1-synMR-TenO1::NatR ade2-1 ura3-1 his3-11 trp1-1 leu2-3 leu2-112 can1-100 |
| LFL001 | Strain harboring the surrogate *SRP54** driven by *pMR* and a deleted native *SRP54*. | MATa HO::pTEF1-synMR-TenO1::NatR Met17::Zif268-SRP54mutcut-TenO1::HygR SRP54△ ade2-1 ura3-1 his3-11 trp1-1 leu2-3 leu2-112 can1-100 |

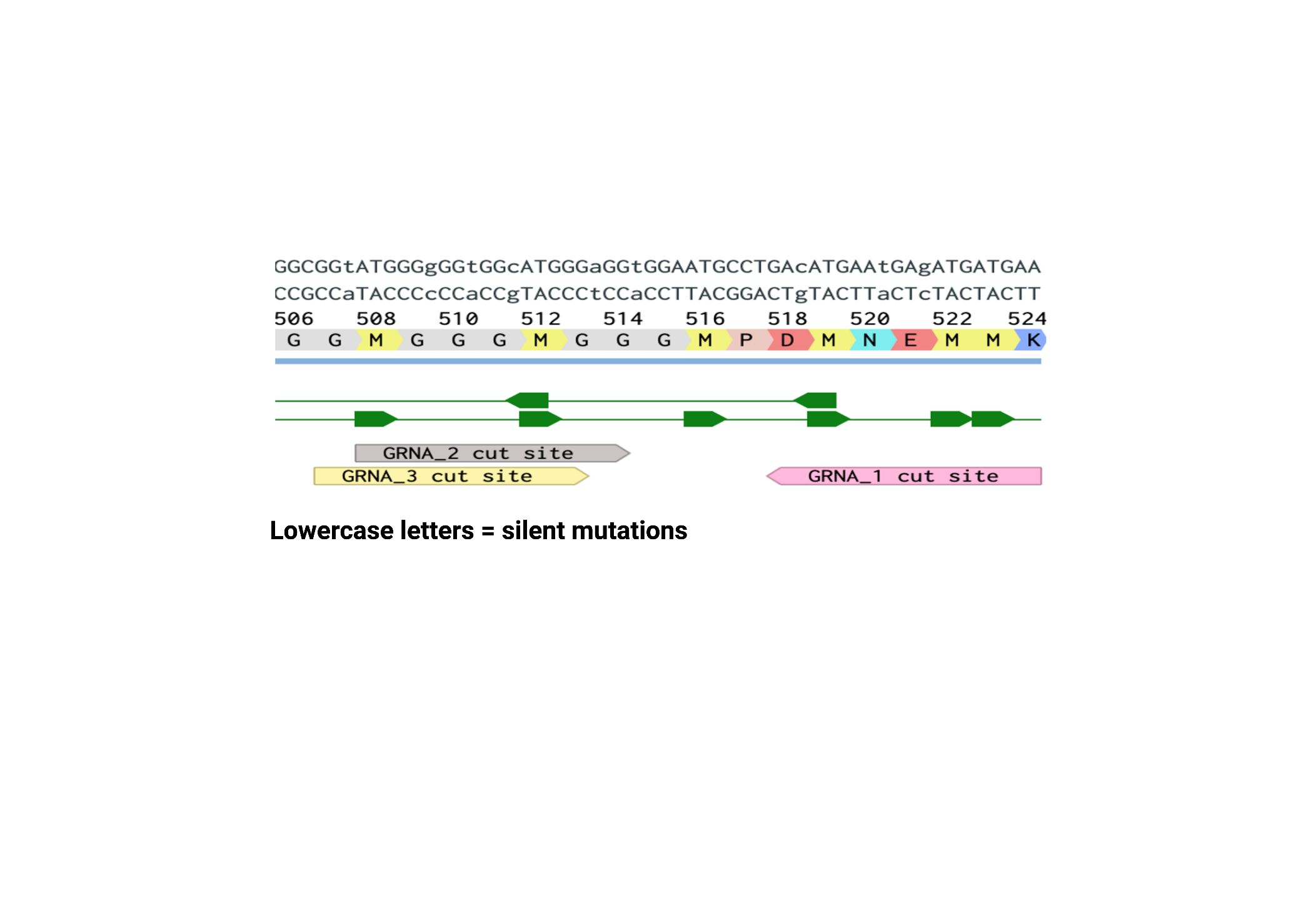

**Supplemental Figure 1: Silent mutations prevent Cas9 from cleaving the newly inserted SRP54***

A series of 9 silent mutations were made to the inserted SRP54 cassette (represented by lowercase letters) to prevent unintentional cleavage of the SRP54* gene by Cas9 during the knockout of the native gene. These mutations were selectively made at gRNA binding sites to create a 3 to 4 base pair discrepancy between the native SRP54 and the newly inserted SRP54*. Aside from these silent mutations, the sequence was left identical to the WT SRP54 to minimize changes in codon usage. A series of 3 guides were tested; however, gRNA 2 was ultimately the one used for the knockout of the native gene.

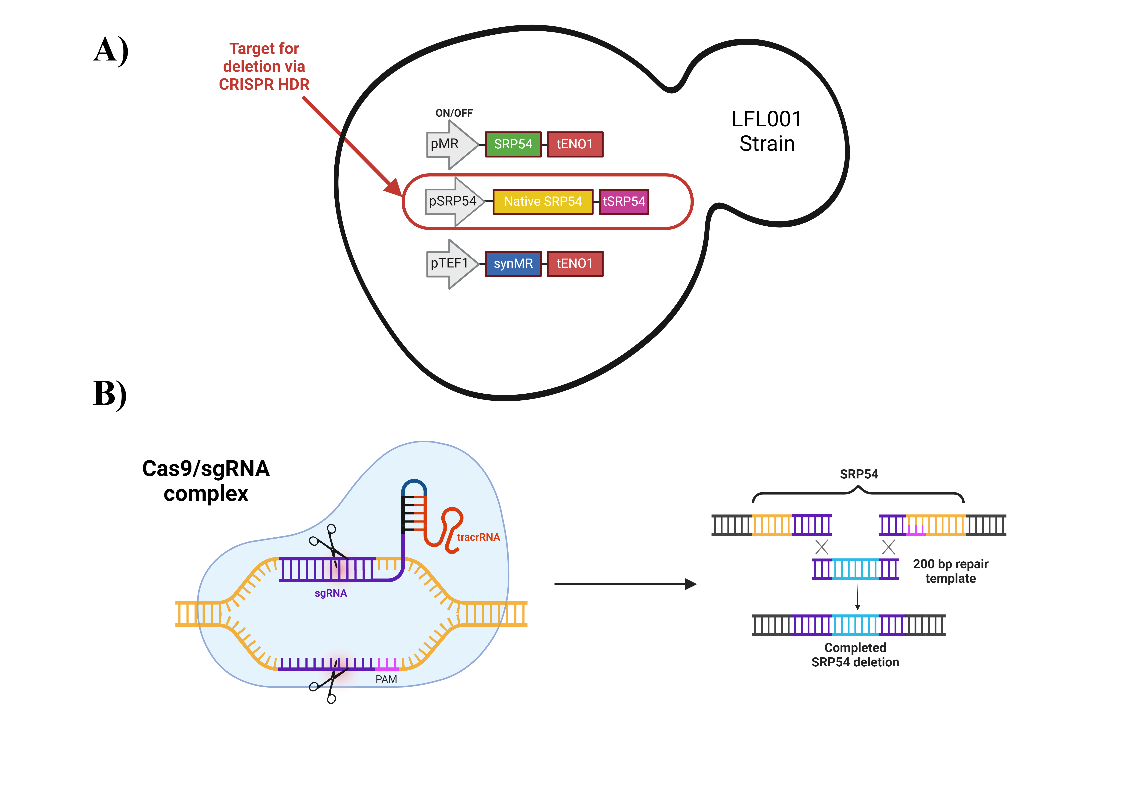

**Supplemental Figure 2:** **Deletion of native *SRP54* via CRISPR HDR**

**A)** After insertion of the *pMR*-driven *SRP54* cassette, the native *SRP54* gene requires removal to create in the final LFL001 strain. **B)** A Cas9-mediated homology-directed repair (HDR) was conducted such that the native *SRP54* open reading frame was replaced with a 200 base pair repair template deleting the gene. Notably, the bi-directional promoter and terminator of *SRP54* were left undisturbed to minimize disruptions to the surrounding regions. A PML107 Cas9 expression vector was used to knock out the native *SRP54* gene. Because *SRP54* is a quasi-essential gene, the *pMR*-driven *SRP54* cassette was toggled "on" by adding 10µM aldosterone to the selection media to promote cell viability during the Cas9-mediated knockout of the native *SRP54* gene. By doing this, a functional copy of SRP54 is expressed throughout the process.

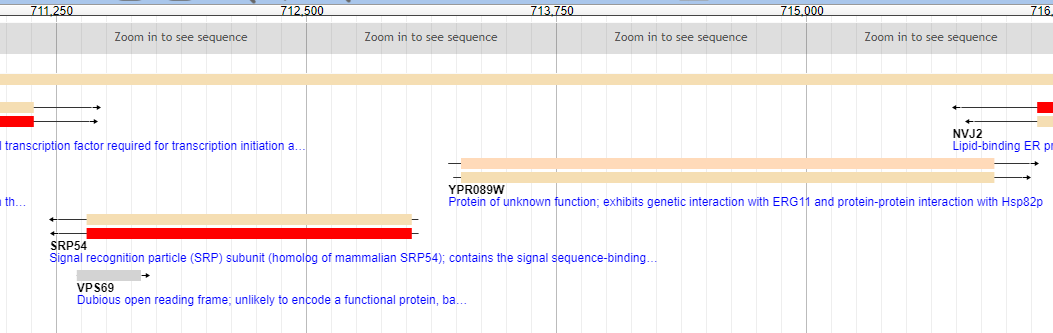

**Supplemental Figure 3: The *SRP54* gene shares its promoter with a protein of unknown function**. Our efforts in designing a suitable repair template involved preserving this promoter region to avoid disrupting the neighboring gene. The repair template thus replaced the open reading frame leaving the promoter and terminator regions undisturbed.

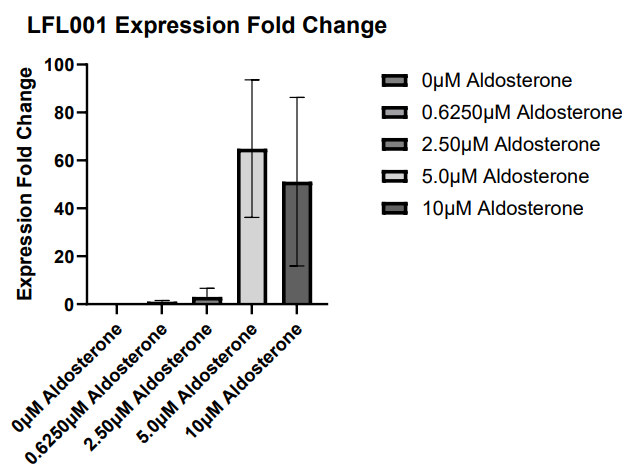

**Supplemental figure 4: LFL001 *SRP54* Expression Fold Change**

*SRP54* expression is dependent upon aldosterone concentration. At aldosterone concentration 0.625µM, the *SRP54* expression at 1.02 expression fold change over the native *SRP54*. At aldosterone concentrations ≥5µM *SRP54* expression levels sharply increase representing extreme overexpression of *SRP54*. Surprisingly, large increases in *SRP54* expression do not result in perceivable differences in the growth rate of the strain. Data represents an N=2 sample size, and error bars represent the calculated standard deviation for each sample.
